# Divergent CPEB prion-like domains reveal different assembly mechanisms for a generic amyloid-like fold

**DOI:** 10.1101/2020.05.19.103804

**Authors:** Rubén Hervás, María del Carmen Fernández-Ramírez, Albert Galera-Prat, Mari Suzuki, Yoshitaka Nagai, Marta Bruix, Margarita Menéndez, Douglas V. Laurents, Mariano Carrión-Vázquez

## Abstract

Functional amyloids are present in a wide variety of organisms ranging from bacteria to humans. Experience-dependent aggregation of the cytoplasmic polyadenylation element-binding (CPEB) prion-like protein to a translationally active state has emerged as a plausible biochemical substrate of long-lasting memories. CPEB aggregation is driven by prion-like domains (PLD) that are highly divergent in sequence across species. Here, we describe the amyloid-like features of the neuronal *Aplysia* CPEB (*Ap*CPEB) PLD *in vitro* using single-molecule and bulk biophysical methods and compare them with those previously reported for neuronal *Drosophila* CPEB, Orb2 PLD. The existence of transient oligomers and mature filaments suggests similarities in the late stages of the assembly pathway for both PLDs. However, while prior to aggregation the Orb2 PLD monomer remains as a random coil in solution, *Ap*CPEB PLD adopts a diversity of conformations comprising *α*-helical structures that evolve to coiled-coil species, suggesting structural differences at the beginning of their amyloid assembly pathways. Our results show how divergent PLDs of CPEB proteins from different species retain the ability to form a generic amyloid-like fold through different assembly mechanisms.

## Main text

New biological traits can emerge from heritable changes in protein-based epigenetic elements known as prions (Alberti et al. 2009; J. C. S. Brown and Lindquist 2009; Volkov et al. 2002; Wickner 1994; Hou et al. 2011), redefining the central dogma in which the heritable information is stored in nucleic acids (Chakravarty and Jarosz 2018). A prion-like protein with a defined role in memory persistence in different species, from *Aplysia* to mammals, is the mRNA-binding cytoplasmic polyadenylation element-binding, CPEB, family of proteins. Discovered first in oocytes as regulators of mRNA translation (Hake and Richter 1994), the aggregated state of neuronal-specific isoforms of CPEB family members was later found to play a key role in long-term synaptic facilitation in *Aplysia* (Si et al. 2010), long-term potentiation of synaptic transmission in mice (Fioriti et al. 2015) and maintenance of long-term memory in both mice and *Drosophila* (Majumdar et al. 2012; Khan et al. 2015; Kruttner et al. 2012; 2015; Keleman et al. 2007; Fioriti et al. 2015; Li et al. 2016; Hervas et al. 2016), through the activation of dormant mRNAs mediated by, at least in *Drosophila*, a self-sustaining amyloid state (Hervas et al. 2020).

Sequence analysis revealed that the folded C-terminus of neuronal CPEB proteins, which comprises two RNA-recognition motifs (RRM) and a ZZ-type zinc finger domain, is highly conserved. By contrast, the N-terminus, which comprises the PLD responsible for the formation of SDS-resistant, functional aggregates of *Ap*CPEB, Orb2 and CPEB3 in the adult brain of *Aplysia, Drosophila*, and vertebrates, respectively (Hervas et al. 2020; Majumdar et al. 2012; Stephan et al. 2015; Si, Lindquist, and Kandel 2003), is highly divergent across species (**Figure 1A, B** and **S1**).

**Figure 1.**
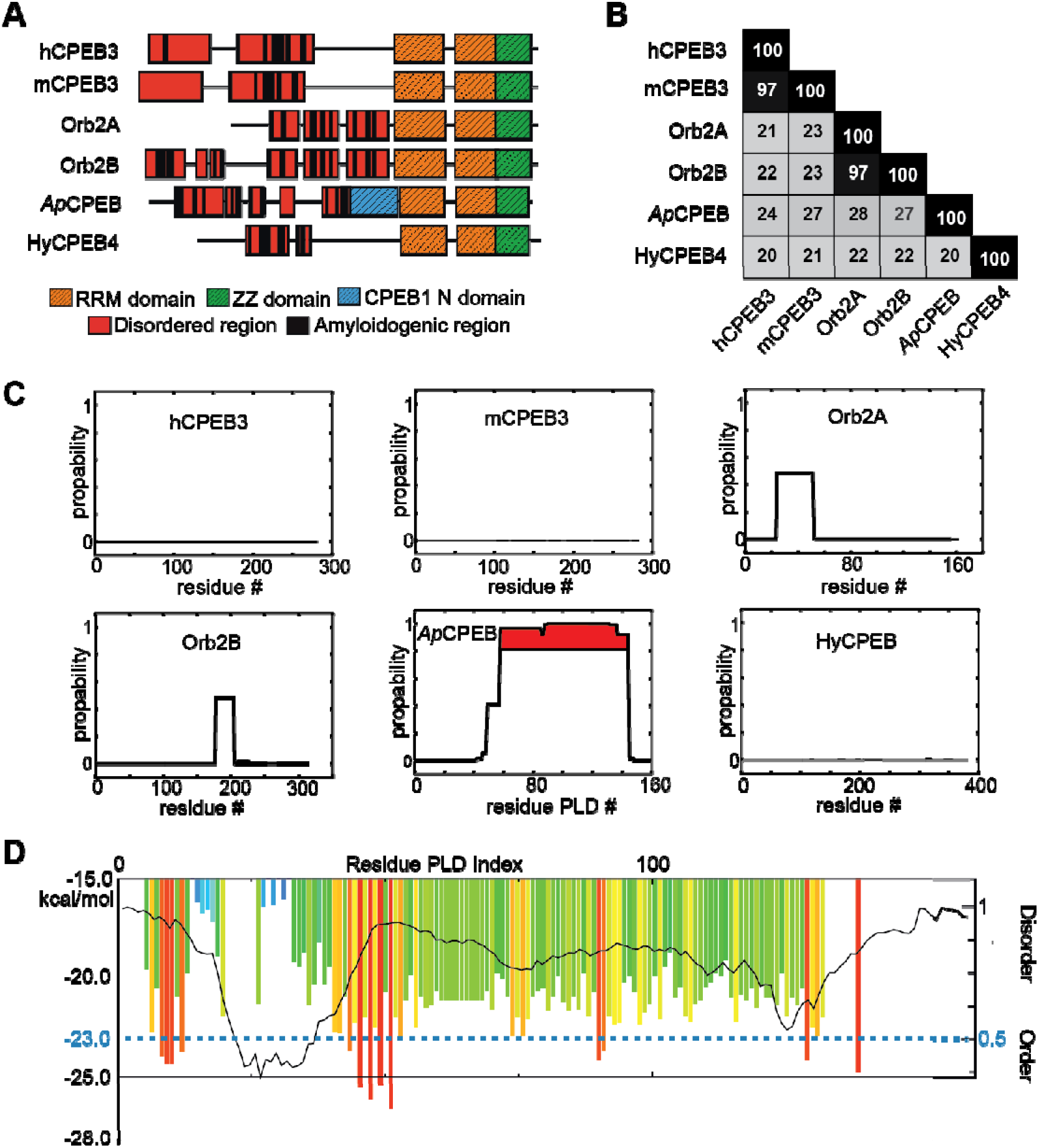
Computational sequence analysis of neuronal CPEB orthologs. **A)** The selected CPEB proteins have a defined role in memory in different species. CPEB4 from *Hydra magnipapillata*, a fresh-water member of the cnidarian, was selected as the most primitive animal to have neurons. The two RRMs (orange) and ZZ domain (green) are located in the highly conserved C-termini. The divergent N-termini of all analyzed CPEB proteins, where the PLD is located, show small amyloid-prone (black boxes) and disordered (red boxes) segments. **B)** Pairwise sequence identity of the N-terminal region represented as a matrix, from the first residue to the first residue of RRM1. *Ap*CPEB PLD and Orb2A PLD share 28% sequence identity. **C)** CC probability (from 0 to 1, *per* residue) obtained using *Coils* algorithm for N-terminal regions of neuronal CPEB isoforms. A 28-residue window was used for the analysis. *Ap*CPEB PLD is the only domain with segments whose CC formation propensity is over 0.8 (highlighted in red), followed by *Drosophila* Orb2A PLD and Orb2B PLD with a propensity near 0.5. **D)** Prediction of disorder propensity using PONDR-FIT, and amyloid-spine formation using ZipperDB, along the 160 residues of the *Ap*CPEB PLD sequence. Dark line shows the predicted disorder score *per* residue. Values above or below 0.5 predict disordered or ordered, respectively. Each colored bar indicates the hexapeptide starting position for amyloid-spine prediction. Segments with lower energy than the threshold at −23 kcal/mol are indicated in red as prone-amyloid segments.

In *Drosophila* nervous system, Orb2 exists in different conformational states. While Orb2 monomers repress synaptic translation, the amyloid state enhances it (Hervas et al. 2020; Khan et al. 2015). Similarly, the neuronal-specific *Ap*CPEB isoform can exist in at least two different conformational states: a soluble form and a *β*-sheet-rich amyloid form with enhanced binding capacity to target mRNAs (Si, Lindquist, and Kandel 2003; Raveendra et al. 2013). In both proteins, the PLD located at the N-terminus plays a key role in the transition to a *β*-sheet-rich state, in spite of their low sequence identity (**Figure 1B** and **S1C**). How divergent PLD sequences drive the assembly of CPEB aggregates in different species to perform a common function remains unclear. To explore the underlying mechanisms during assembly, we have characterized specific structural features of *Ap*CPEB PLD, from the monomer to the aggregated state, and compare them with those of Orb2A (Hervas et al. 2016). The role of coiled-coils (CCs) in aggregation and activity of Q/N-rich proteins *in vivo* has been reported (Fiumara et al. 2010). By using the Coils algorithm (Lupas, Van Dyke, and Stock 1991), we found that *Ap*CPEB PLD scored as the highest predicted hit to form CC domains (**Figure 1C**). As shown in **Figure 1D**, its sequence contains regions prone to be disordered and form amyloid spines. The next predicted hit to form CC corresponded to Orb2 PLD, which was previously reported, by Circular Dichroism (CD), to form a random coil conformation in solution (Hervas et al. 2016). The CC propensity of the mouse, human and hydra orthologs (mCPEB3, hCPEB3 and hyCPEB PLDs) was insignificant (**Figure 1C**).

Compatible with CC prediction, *Ap*CPEB PLD showed a dominant *α*-helical spectrum by CD (**Figure 2A**), similar to full-length *Ap*CPEB (Raveendra et al. 2013). After 72h incubation, the CD signal decreased to approximately 10% of the initial value (**Figure 2A**). Q-binding peptide 1 (QBP1), a peptide known to block the *β*-structure transition and the aggregation of poly-Q containing proteins (Nagai et al. 2007; Hervas et al. 2016; Ramos-Martin et al. 2014; Hervas et al. 2012), and a scrambled version of it (SCR, (Tomita et al. 2009)), were used to determine the nature of the loss of signal. *Ap*CPEB PLD directly interacted with QBP1 through an exothermic reaction, as revealed by isothermal titration calorimetry (ITC). By contrast, SCR did not show significant interaction, according to the net heat exchange (**Figure S2**), suggesting that the observed loss of CD signal is, at least in part, likely due to *Ap*CPEB PLD aggregation over time and that QBP1 was able to lower it (**Figure 2A**).

**Figure 2.**
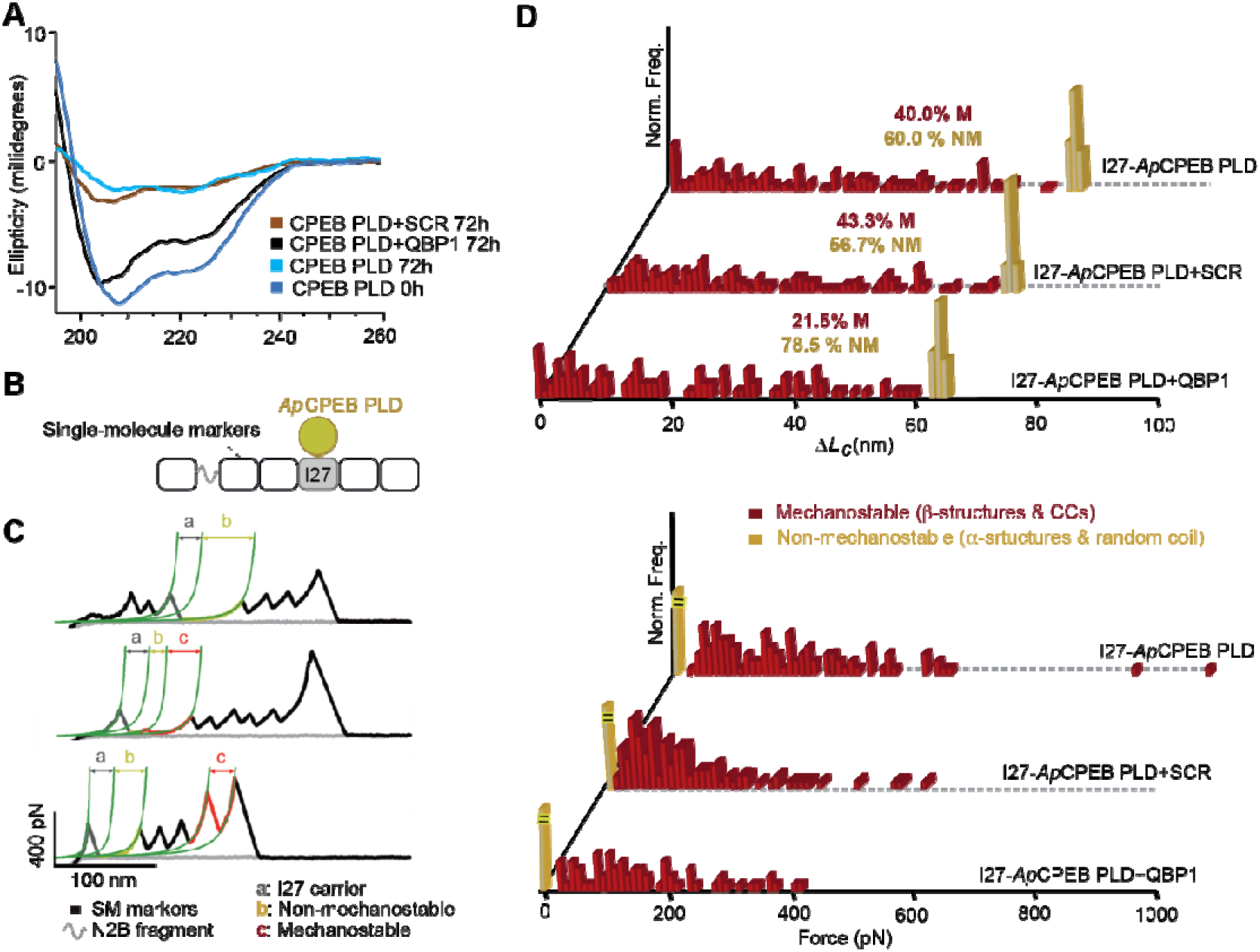
Structural characterization of *Ap*CPEB PLD monomer. **A)** The far-UV CD spectra shows two minima at ~222 nm and 208 nm and a maximum below 200 nm, which are characteristic of *α*-helix-rich conformations. In the presence of QBP1, but not SCR peptide, the loss of signal intensity over time is reduced. **B)** Schematic cartoon of pFS-2 vector using I27 as a carrier (gray) and hosting *Ap*CPEB PLD (yellow). Singlemolecule (SM) markers are represented in black. The polyprotein encoded by pFS-2 also contains a random coil region (a fragment of titin N2B, grey line) that acts as a spacer to overcome the noisy proximal region of the force-extension recordings. **C)** Representative force-extensions recordings. Traces in yellow correspond to NM events. M events, in red, show different mechanical stability (*F*_u_) and increased contour length (Δ*L_c_*) values. “b”+”c” values complete the length of the fully stretched *Ap*CPEB PLD. **D)** Δ*L_c_* and *F*_u_ histograms for polyproteins carrying *Ap*CPEB PLD reveal a rich conformational polymorphism in terms of Δ*L_c_* (top panel) and *F* (bottom panel), ranging from NM events (yellow bars) to M events (red bars, *n*=145). In the presence of a 1:10 molar excess of QBP1, the mechanical conformational polymorphism is lowered, while SCR has no apparent effect (QBP1 *n*= 186, SCR *n*= 144).

Next, we used single-molecule force spectroscopy based on atomic force microscopy (AFM-SMFS) using the length-clamp mode together with a protein engineering strategy, which was previously used to analyze conformational diversity of amyloid-forming proteins at the monomer level (Hervas et al. 2012; 2016; Fernandez-Ramirez et al. 2018). Here, *Ap*CPEB PLD is mechanically protected inside a carrier protein (I27 module from human cardiac titin) by cloning its code into the pFS-2 plasmid (**Figure 2B** and **S3A**) (Oroz, Hervas, and Carrion-Vazquez 2012). Upon *Ap*CPEB PLD insertion, 2D NMR spectroscopy showed that I27 moiety retains its native tertiary structure (**Figure S3B**). AFM-SMFS showed that the monomeric *Ap*CPEB PLD fluctuates over a wide conformational space that includes two main populations: 60.0% of non-mechanically-resistant (NM; unfolding force, *F*, ≤20 pN) and 40.0% of mechanically-resistant (M; *F*>20 pN), spanning a wide range of stabilities (**Figure 2C, D** and **Supplemental Table 1**). M recordings are mainly attributed to the unraveling of the *β*-structures (Oberhauser and Carrion-Vazquez 2008), although low forces, still detected by AFM-SMFS, can be due to coiled coil mechanical unfolding (A. E. X. Brown et al. 2007; Goktas et al. 2018; Bornschlogl and Rief 2008; Schwaiger et al. 2002). By contrast, NM events could originate either from RC or isolated *α*-helix conformers (Oberhauser and Carrion-Vazquez 2008). Similar results were obtained using ubiquitin (Ubi) as carrier (Oroz, Hervas, and Carrion-Vazquez 2012), ruling out potential artefactual effects due to the use of a specific carrier (**Figure S4A**). Finally, QBP1 lowered the formation of M conformers, while neither SCR (**Figure 2D**) nor DMSO (used as QBP1/SRC vehicle, **Figure S4B**) had an effect. In any case, formation of preferential conformers was not observed (**Figure S4C and Supplemental Table 1**).

Far-UV CD spectra deconvolution of freshly purified *Ap*CPEB PLD revealed that a great proportion of conformations are structured (**Figure 3A**), in contrast to its ortholog Orb2A PLD, which remains as a random coil (Hervas et al. 2016). In addition, primary structural changes associated with protein assembly can be monitored by far-UV CD. The gradual loss of signal due to protein aggregation is accompanied by an increase in the 220/208 ellipticity ratio (**Figure 3B**), which has been associated to the presence of CCs (Lilliu et al. 2018). Concentration-normalized spectra at 72 h do not overlap, suggesting that the observed differences are due to structural rearrangements over time (**Figure 3C**). This observation is consistent with an aggregation mechanism whose early steps would involve a transition from isolated *α*-helix to CC (Fiumara et al. 2010; M. Chen, Zheng, and Wolynes 2016). Accompanying the CCs formation during the first 72 h, oligomeric SDS-sensitive species, recognized by the A11 conformational antibody, also were formed (Kayed et al. 2003), which evolved rapidly to SDS-resistant OC-reactive species (Kayed et al. 2007) (**Figure 3D**). OC-reactive species showed amyloid-like features, such as Congo Red binding (**Figure 3E**) and the formation of annular assemblies associated to unbranched filaments of different lengths and approximately 7-25 nm wide (**Figure 3F**). QBP1, as shown previously by CD (**Figure 2A**), reduced *Ap*CPEB PLD aggregation (**Figure 3D**).

**Figure 3.**
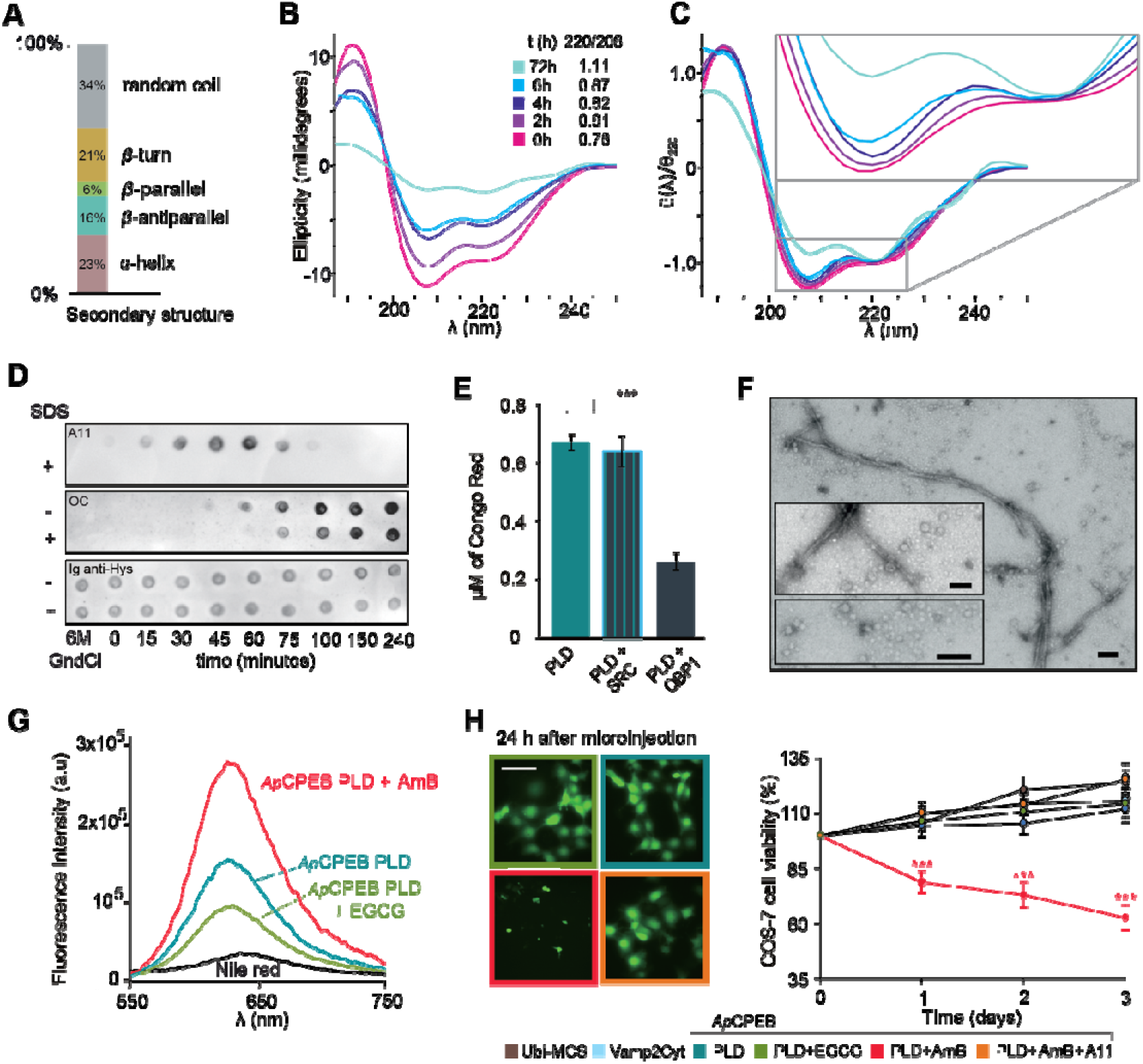
Dissection of *Ap*CPEB PLD amyloid-like assembly pathway. **A)** Average secondary structure content of soluble *Ap*CPEB PLD conformers derived by far-UV CD spectrum deconvolution. The fractions of *α*-helix and RC conformers that likely originate the NM events, as well as those of containing *β*-structures (likely M events) are estimated. **B)** Raw far-UV CD spectra of *Ap*CPEB PLD over time (left). At zero time, the spectrum shows minima at 220 and 208 nm and a maximum at around 195 nm, characteristic of *α*-helical structure. A loss of signal with time is accompanying an increase in the 220/208 ratio, which is associated to CC formation. **C)** The change of shape with time can be better appreciated when the relative intensities of the spectra, with relation to the respective value at 220 nm, are compared to compensate the loss of signal. Normalized spectra do not overlap, particularly at 72h, suggesting the occurrence of structural changes over time. **D)** SDS-sensitive A11-reactive oligomeric species appeared from ~15 to 75h and evolved to SDS-resistant OC-reactive aggregates, presumably amyloid-like, beyond that time point. **E)** The concentration (*μ*M) of Congo red bound to *Ap*CPEB PLD amyloid-like species was lowered in the presence of QBP1. Data is represented as mean ± SEM: \*\*\**p*<0.001 (One-way ANOVA and Tukey posttest). **F)** Representative electron micrographs of oligomers and amyloid-like filaments are shown in a general view and magnified images. Scale bar: 50 nm. **G)** Nile red fluorescence emission spectra of *Ap*CPEB PLD, *Ap*CPEB PLD:AmB and *Ap*CPEB PLD:EGCG complexes. While treatment with AmB increases the Nile red fluorescence values, EGCG causes the opposite effect. **H)** *Ap*CPEB PLD microinjection in the COS-7 cell line. Left panel: fluorescence micrographs of COS-7 cells 24h after the microinjection of indicated samples along with fluorescein-labelled dextran. *Ap*CPEB PLD:AmB microinjection decreased the number of live cells after 24h. Scale bars: 100 *μ*m. Right panel: survival curves of COS-7 microinjected cells. None of the samples, but the *Ap*CPEB PLD species trapped with AmB, cause toxicity. The toxic phenotype was rescued by the addition of A11 antibody. Data are represented as the mean ± SEM: \*\*\**p*<0.00 (red asterisks) *Ap*CPEB PLD:AmB *versus* remaining samples; Two-way ANOVA and Bonferroni post-test. Ubi-MCS and the cytoplasmic region of VAMP2 are used as controls of folded and disordered proteins, respectively. The number of cells microinjected per sample was *n* = 100-200.

Formation of A11-reactive species that rapidly evolved to a mature filament state with amyloid-like features was also found in Orb2A PLD (Hervas et al. 2016). Small-molecule inhibitors Amphotericin B (AmB) and (-)-epigallocatechin gallate (EGCG) interfered with *Ap*CPEB assembly (**Figure S5**), as shown for other amyloids (Krishnan et al. 2012; Hervas et al. 2016). Thus, *Ap*CPEB PLD was trapped in an A11-reactive conformation by AmB and in an OC-reactive conformation by EGCG (**Figure S5**). Oligomers trapped in an A11-reactive state exposed more hydrophobic residues to the solvent (**Figure 3G**), and induced cell death upon microinjection into COS-7 cells, contrary to OC-reactive species or untreated *Ap*CPEB PLD (**Figure 3H**). Oligomers trapped in the A11-reactive state incubated with A11 antibody (complex:A11, 100:1) neutralized its toxic phenotype (**Figure 3H**), suggesting that exposure of hydrophobic residues in aggregates is associated with cell toxicity (Campioni et al. 2010; S. W. Chen et al. 2015; Liu et al. 2012).

These results indicate that the amyloid assembly of *Drosophila* and *Aplysia* CPEB PLD orthologs converges at the oligomeric state. The presence of similar shortlived A11-reactive oligomers that evolve rapidly to OC- and CR-reactive amyloid filaments supports the idea that toxic A11 reactive oligomers may be intrinsically ephemeral in functional amyloids (Krishnan et al. 2012; Hervas et al. 2016). However, we found key differences at the start of each assembly pathway, likely related to their divergent PLD sequences. Both CPEB PLDs display a broad conformational polymorphism at the monomer level measured by AFM-SMFS, reinforcing the notion that monomers of amyloid-forming proteins rapidly fluctuate between a wide conformational space (Mukhopadhyay et al. 2007; Ferreon et al. 2010). However, contrary to Orb2A PLD (Hervas et al. 2016), *Ap*CPEB PLD is structured in solution and, *in vitro*, early aggregation is likely mediated by the formation of CCs, in agreement with previous observations for other Q/N-rich proteins (Fiumara et al. 2010). In this model, CCs correspond to an alternate stable oligomeric intermediate that would allow aggregation to be regulated (Raveendra et al. 2013; M. Chen, Zheng, and Wolynes 2016). Once formed, CCs may represent intermediate assembly structures mediating the conformational transition of neighboring random coil segments, or facilitating their own conversion, into *β*-sheet multimers. (Fiumara et al. 2010; M. Chen, Zheng, and Wolynes 2016) (**Figure S6**).

Taking together, our findings show that the PLD of neuronal CPEB proteins from different species, in spite of their highly divergent sequence, has evolved to retain the ability to form an amyloid-like fold through different assembly mechanisms. It has been reported that PLDs from different proteins are functionally interchangeable (Hervas et al. 2016; Si, Lindquist, and Kandel 2003), suggesting that attaining the amyloid-like fold may be sufficient to preserve the protein function. Moreover, their differential amyloidogenesis at the early stages may also indicate the presence of different regulation systems, an issue that should be addressed in the future. Those future studies should answer the question of whether this structural diversity at the early stages of CPEB assembly has a biological role or if, on the contrary, it is just an epiphenomenon due to the lack of structural evolutionary constraints of their unstructured regions, which could have resulted in biologically equivalent sequences.

## Acknowledgements

We thank to M. I. Maher (EM facility, Cajal Institute) for technical assistance and to J. M. Valpuesta for the access to his glow-discharge apparatus. The work was funded by two joint grants from the Ministry of Economy and Competitiveness to MCV (SAF2013-49179-C2-1-R and SAF2016-76678-C2-1-R) and DVL (SAF2013-49179-C2-2-R and SAF2016-76678-C2-2-R) and grants of the Ministry of Economy and Science (BFU2015-70072-R) and the CIBER de Enfermedades Respiratorias (CIBERES; ISCIII) to MM.

## Author contributions

Conceptualization, MCV and RH; investigation, RH, MCFR, AGP, MS, YN, MB, MM, DVL and MCV; resources, YN, MB, MM, DVL and MCV; supervision, MCV; writing, original draft, RH and MCFR; writing–review and editing, all authors. Competing interests: the authors declare no competing financial interests.

## Materials and Methods

### Cloning

All constructs containing the N/Q-rich region of the neuronal-specific *Aplysia* CPEB (residues 1-160) were cloned by PCR using a full-length clone (Addgene) as the template. The oligonucleotides used are listed in the **Supplemental Table 2**. All sequences were first cloned into the pCR2.1 (Invitrogen) or pT7Blue (Novagen) vectors, verified, and subcloned, using the *Escherichia coli* strains DH5*α* (Invitrogen) and XL1-Blue (Stratagene), into i) the pFS-2 vector, using the AgeI-SmaI restriction sites of the multicloning site (MCS) region in the I27 or Ubi carrier module (Oroz, Hervas, and Carrion-Vazquez 2012) for AFM-SMFS analysis; or ii) into the pET28a vector (Novagen), using the NheI and XhoI restriction sites, to construct *Ap*CPEB PLD and *Ap*CPEB PLD-fusion proteins for all the remaining experiments, as in (Oroz, Hervas, and Carrion-Vazquez 2012). Finally, all sequences cloned in expression vectors were verified by sequencing both DNA strands.

### Protein Expression

All isolated, carrier-guest monomers and pFS-2 fusion proteins were expressed in the *E. coli* C41(DE3) or BL21(DE3) strains (Invitrogen). Bacterial cultures were grown at 37°C until they reached an OD_595_ of 0.4-1, and then expression was induced by addition of 1 mM IPTG for 4 h. Then, they were lysed using lysozyme and sonication pulses. The recombinant proteins were purified by Ni^2+^-affinity chromatography using Histrap HP FPLC columns (GE Healthcare) and a size-exclusion chromatography using a HiLoad 16/60 200 PG column (GE Healthcare) on an FPLC apparatus (ÄKTA Purifier, GE Healthcare). Protein concentration was determined by absorbance at 280 nm using the molar extinction coefficient of each protein.

### Computational analysis

Sequences were obtained from Uniprot and GenBank and analyzed in PFAM (Finn et al. 2016) and InterPro (Mitchell et al. 2015) to study the presence of conserved domains. Sequences were aligned using Clustal Omega with default parameters and represented using boxshade. The pairwise identity as calculated by Clustal was represented as identity matrices. The N-terminal region was selected as the region from the N-terminus to the first RRM motif; RRM1 and RRM2 sequences were those identified by Interpro, while ZZ were identified by PFAM.

We used the ZipperDB database to identify sequence segments with tendency to form the steric zipper spines of amyloid filaments (Goldschmidt et al. 2010) (https://services.mbi.ucla.edu/zipperdb/). Order/disorder prediction was made using a consensus artificial neural network prediction method, Predictor Of Naturally Disordered Regions PONDR-FIT (Molecular Kinetics). This metapredictor was developed by combining the outputs of several individual disorder predictors (PONDR-VLXT, PONDR-VSL2, PONDR-VL3, FoldIndex, IUPred, and TopIDP) (Xue et al. 2010).

### NMR

2D ^1^H NOESY NMR spectra were measured using samples at 1 mM in 10 mM KH_2_PO_4_, pH 4.7 with 10% D_2_O and at 25 °C, using a Bruker AV 800 spectrometer (Bruker BioSpin) equipped with a ^1^H, ^13^C, ^15^N cryoprobe and Z-gradients. The temperature was calibrated using an ethanol sample and the signal of the trimethyl moiety of sodium 2,2-dimethyl-2-silapentane-5-sulfonate (DSS) was used as an internal reference of the chemical shift. Selective pre-s aturati on or a WATERGATE module (Piotto, Saudek, and Sklenar 1992) were used to reduce the signal of the water present in the sample. The spectra were analyzed using TopSpin 2.0 (Bruker BioSpin). The assigned signals in the I27 spectra were based on previously reported data (Improta, Politou, and Pastore 1996).

### CD

Far-UV CD spectra of *Ap*CPEB PLD samples, at a concentration of 2 *μ*M in 10 mM KH_2_PO_4_ pH 4.7, were recorded using a JASCO-J810 spectropolarimeter (JASCO Inc.) equipped with a Peltier temperature control unit and using quartz cuvettes of 1 mm cellpath length. When required, after correction by subtraction of the buffer contribution, the spectra were converted into molar ellipticity ([Θ]), using the average molecular masses *per* residue with Spectra Manager software (Jasco Inc.). The secondary structure content was quantified by CD spectra deconvolution using the CDNN analysis program (Bohm, Muhr, and Jaenicke 1992). Once the first protein spectrum (hour 0) was recorded, the protein was incubated at 37.0°C without stirring (with 0.02% NaN_3_), and additional spectra were collected over the following hours. For comparative purposes, and to minimize the impact of signal loss, the spectra registered as a function of time were normalized by dividing full trace intensities by its own value acquired at 220 nm, where is the lower recorded value in the second minimum of the spectra. The concentration of QBP1 and SCR (10 *μ*M with a final DMSO concentration below 0.01%, to avoid DMSO interference with the ellipticity (Θ) measurements) was 1:5 in molar excess. QBP1-M8 (Ac-WKWWPGIF-NH_2_) and SCR (Ac-WPIWKGWF-NH_2_) peptides with the N- and C-termini acetylated and amidated, respectively, were synthesized at the Proteomic Facility of the CBMSO/CSIC-UAM using solid-state Fmoc chemistry with N-terminal acetylation and C-terminal amidation.

### ITC

ITC experiments to examine the interaction of *Ap*CPEB PLD with the minimal active core of the QBP1 (Ac-WKWWPGIF-NH_2_) and SCR (Ac-WPIWKGWF-NH_2_) peptides were carried out in a VP-ITC microcalorimeter at 25°C. The proteins were equilibrated in PBS pH 7.4 by size exclusion chromatography, and the equilibration buffer was used to prepare the peptide solutions. Protein/peptide binding was tested by successive injections of the protein (10 to 25 *μ*l each) into the reaction cell loaded with peptide at a high final (peptide)/(protein) molar ratio. The apparent heat of reaction for each injection was obtained by integration of the peak area. The heat developed with the protein or peptide dilutions was determined in separate runs, loading either the sample cell or the injection syringe with buffer in the conditions used for the binding experiments. The complexity of the system and the lack of precise information on the distribution of the *Ap*CPEB PLD conformations before and after complex formation precluded the quantitative analysis of the titration curves. The protein (*Ap*CPEB PLD, 550 *μ*M) and ligand (QBP1 and SCR, 75 *μ*M and 51 *μ*M, respectively) concentrations in the loading solutions were measured spectrophotometrically using their respective extinction coefficients.

### CR binding assay

Congo Red binding assays were performed in 10 *μ*M *Ap*CPEB PLD samples that were dialyzed in PBS, pH 7.0. Both SCR and QBP1 peptides were used at a concentration of 50 *μ*M. Before the analysis, samples were aged for 10 days at 37.0°C without stirring in the presence of 0.02% NaN_3_. After the incubation time, the protein solutions were mixed with a 30 *μ*M Congo Red solution (in 5 mM sodium phosphate buffer+300 mM NaCl, pH 7.5) and incubated at room temperature for 30 min. Congo Red binding was calculated by measuring bathochromic and hyperchromic shifts in the samples, using a UV-visible spectrophotometer (Nanodrop, Thermo Scientific), and applying the following equation Congo Red *μ*/mol/l) = A_540_/25,295 – A_480_/46,306 (Wurth, Guimard, and Hecht 2002). Congo Red bound to amyloid aggregates exhibited typical apple-green birefringence, as monitored under polarized light Leica DMI6000B inverted optical microscope (Leica Microsystems).

### Dot Blot analysis

To test if *Ap*CPEB PLD (isolated and fused in the I27 carrier) forms A11 and OC-reactive species, *Ap*CPEB PLD aliquots at a concentration of 10 *μ*M in PBS at pH 7.0 were used. For EGCG and AmB immune-dot blot analysis, *Ap*CPEB PLD in monomeric state was prepared by denaturing it in 6M GndCl (considered as the starting time of the reaction) and then diluting it in 5 mM potassium phosphate + 150 mM NaCl, pH 7.4 to a final protein concentration of 2.5-5 *μ*M. DMSO (AmB vehicle) or 4:1 molar excess of AmB and EGCG was added. In all the cases, 2 *μ*l of sample were spotted onto a nitrocellulose membrane. After blocking the membrane for 1h at room temperature with 10% non-fat milk in TBS containing 0.01% Tween-20, the membrane was incubated at room temperature for 1 h with the polyclonal specific anti-oligomer A11 antibody (Life Technologies) or the fibril-specific monoclonal antibody OC (Millipore), diluted to 1:1,000 in 3% BSA TBS-T. The membranes were washed for 5 min three times with TBS-T before incubating at room temperature for 1 h with the secondary antibody (anti-rabbit HRP conjugated anti-rabbit IgG [GE Healthcare] diluted 1:5,000 in 3% BSA/ TBS-T). After washing the membranes three times in TBS-T buffer, the blots were developed with ECL Plus chemiluminescence kit from Amersham-Pharmacia (GE Healthcare).

### TEM imaging

10 *μ*M *Ap*CPEB PLD protein were dialyzed in PBS, pH 7.0 for all TEM measurements. 10 *μ*l of sample were adsorbed onto carbon-coated 300-mesh copper grids (Ted Pella) and negatively stained for 30 s, using 1-2% uranyl acetate. Immediately before use, the carbon-coated grids were *glow-discharged* to enhance their hydrophilicity using an Emitech K100X apparatus (Quorum Technologies). Images were taken on a JEOL 1200EX II (Jeol Limited) electron microscope equipped with a CCD Megaview III camera (Olympus Soft Imaging) at an acceleration voltage of 80 kV.

### AFM-SMFS

Double-blind SMFS experiments were performed using 2 *μ*M of the pFS-2+*Ap*CPEB PLD polyprotein in 10 mM Tris-HCl, pH 7.5 at room temperature. Proteins were kept at 4°C between sessions and NTA-Ni^2+^ functionalized coverslips were used as a substrate to attach the pFS-2 polyprotein through the His-tag present at their N-terminus (Oroz, Hervas, and Carrion-Vazquez 2012). The details of the stringent criteria used for selecting *bona fide Ap*CPEB PLD single-molecule recordings can be found in (Hervas et al. 2012). For experiments performed in the presence of peptides, we incubated the protein sample with QBP1 (20 *μ*M in DMSO) and SCR (20 *μ*M in DMSO), and the mixed samples were incubated overnight at 4°C before performing the measurements. The custom-made single-molecule AFM used and its operation mode were described previously (Valbuena et al. 2009). Before each experiment, the cantilever tip was cleaned for 5 min using a UV lamp (UV/Ozone ProCleaner™ Plus, Bioforce Nanosciences Inc.). The spring constant of each individual Si_3_N_4_ cantilever (MLCT-AUNM, Veeco Metrology Group; and Biolever, Olympus) was calculated using the equipartition theorem (Florin et al. 1995). Experiments were performed at a constant pulling speed of 0.4 nm/ms in the length-clamp mode (Carrión-Vázquez et al. 2007). The data were analyzed using Igor Pro 6 (Wavemetrics) and the WLC model of polymer elasticity (Bustamante et al. 2004):

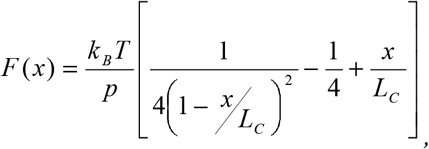

where *F* is the stretching force, *p* is the persistence length, *x* is the end-to-end length, and *L_c_* is the contour length of the stretched protein. *Lc* and *p* are the adjustable parameters.

### Experiments with AmB and EGCG

For the experiments performed in the presence of small molecules, the protein samples with AmB (8 *μ*M in DMSO) and EGCG (20 *μ*M in 10 mM Tris-HCl, pH 7.5) were incubated overnight at 4°C before performing the measurements.

### Nile Red binding assay

Samples of *Ap*CPEB PLD (5 *μ*M in PBS, pH 7.0) in the presence or in absence of 4:1 molar excess of AmB and EGCG were removed at 100 min of incubation at 37°C. A three-fold excess of Nile Red was added and the fluorescence spectrum was recorded using a Jobin Yvon Horiba FluoroMax 4 instrument (Horiba Jobin Yvon) and a 3 mm square quartz cuvette at 25.0 °C. Spectra were recorded in an emission wavelength range of 550-750 nm, with a 120 nm/min scan speed. Spectra of AmB and EGCG alone were also recorded and subtracted from the spectrum of the *Ap*CPEB PLD complex. The excitation and emission wavelengths were calibrated using a fine Xe emission line and the Raman water peak, respectively.

### Single-cell protein microinjection

The COS-7 cell line was grown and maintained in supplemented DMEM with 10% (v/v) FBS. The day before performing the microinjection, cells were plated on a 35-mm dish at a density of 1×10^5^ cells/dish. The day of, and just before the single-cell microinjection experiment, the *Ap*CPEB PLD protein, at 2.5 *μ*M in PBS 7.4, was incubated in the presence or absence of four-fold molar excess of AmB and EGCG for 100 min at 37.0°C to allow complex formation. *Ap*CPEB PLD:AmB:A11 ternary complex was formed by incubation for 3h at room temperature of the previously formed *Ap*CPEB PLD:AmB with the A11 antibody (100:1, binary complex:A11). The samples were double-blinded microinjected into the cytoplasm of single COS-7 cells (n=100-200 cells per sample) along with fluorescein-labeled dextran (molecular weight 10000, Life Technologies) using a micromanipulator (Narishige). The experiments were carried out in a double-blind manner and were repeated three times for each sample. Fluorescence micrographs were acquired using a CCD camera: model C4742-95-12ER (Hamamatsu Photonics). Ubi-MCS and VAMP2Cyt were microinjected as negative controls. Cell viability was monitored by counting the number of fluorescein-positive cells 3 h after microinjection under an IX70 fluorescence microscope (Olympus), and this value was assigned as 100% cell viability. During the three following days, we counted the number of fluorescein-positive cells every 24 h to calculate the cell survival rate. Data are represented as mean ± s.e.m. Two-way ANOVA and Bonferroni post-test and One-way ANOVA and Tukey post-test were used as statistical methods for timecourse survival curves and survival rates, respectively, after 24 h. Statistical analyses were performed using GraphPad Prism 5 (GraphPad Software).

## References

Alberti, Simon, Randal Halfmann, Oliver King, Atul Kapila, and Susan Lindquist. 2009. “A Systematic Survey Identifies Prions and Illuminates Sequence Features of Prionogenic Proteins.” Cell 137 (1): 146–58. https://doi.org/10.1016/j.cell.2009.02.044.

Bohm, G, R Muhr, and R Jaenicke. 1992. “Quantitative Analysis of Protein Far UV Circular Dichroism Spectra by Neural Networks.” Protein Engineering 5 (3): 191–95.

Bornschlogl, Thomas, and Matthias Rief. 2008. “Single-Molecule Dynamics of Mechanical Coiled-Coil Unzipping.” Langmuir □: The ACS Journal of Surfaces and Colloids 24 (4): 1338–42. https://doi.org/10.1021/la7023567.

Brown, André E.X., Rustem I. Litvinov, Dennis E. Discher, and John W. Weisel. 2007. “Forced Unfolding of Coiled-Coils in Fibrinogen by Single-Molecule AFM.” Biophysical Journal. https://doi.org/10.1529/biophysj.106.101261.

Brown, Jessica C S, and Susan Lindquist. 2009. “A Heritable Switch in Carbon Source Utilization Driven by an Unusual Yeast Prion.” Genes & Development 23 (19): 2320–32. https://doi.org/10.1101/gad.1839109.

Bustamante, Carlos, Yann R Chemla, Nancy R Forde, and David Izhaky. 2004. “Mechanical Processes in Biochemistry.” Annual Review of Biochemistry 73: 705–48. https://doi.org/10.1146/annurev.biochem.72.121801.161542.

Campioni, Silvia, Benedetta Mannini, Mariagioia Zampagni, Anna Pensalfini, Claudia Parrini, Elisa Evangelisti, Annalisa Relini, et al. 2010. “A Causative Link between the Structure of Aberrant Protein Oligomers and Their Toxicity.” Nature Chemical Biology 6 (2): 140–47. https://doi.org/10.1038/nchembio.283.

Carrión-Vázquez, Mariano, Andres Oberhauser, Hector Diez, Rubén Hervás, Javier Oroz, Jesús Fernández, and David Martínez-Martín. 2007. “Protein Nanomechanics - as Studied by AFM Single-Molecule Force Spectroscopy.” In Advanced Techniques in Biophysics, 163–245. https://doi.org/10.1007/3-540-30786-9_8.

Chakravarty, Anupam K, and Daniel F Jarosz. 2018. “More than Just a Phase: Prions at the Crossroads of Epigenetic Inheritance and Evolutionary Change.” Journal of Molecular Biology 430 (23): 4607–18. https://doi.org/10.1016/j.jmb.2018.07.017.

Chen, Mingchen, Weihua Zheng, and Peter G Wolynes. 2016. “Energy Landscapes of a Mechanical Prion and Their Implications for the Molecular Mechanism of LongTerm Memory.” Proceedings of the National Academy of Sciences of the United States of America 113 (18): 5006–11. https://doi.org/10.1073/pnas.1602702113.

Chen, Serene W, Srdja Drakulic, Emma Deas, Myriam Ouberai, Francesco A Aprile, Rocio Arranz, Samuel Ness, et al. 2015. “Structural Characterization of Toxic Oligomers That Are Kinetically Trapped during Alpha-Synuclein Fibril Formation.” Proceedings of the National Academy of Sciences of the United States of America 112 (16): E1994–2003. https://doi.org/10.1073/pnas.1421204112.

Fernandez-Ramirez, Maria Del Carmen, Ruben Hervas, Albert Galera-Prat, Douglas V Laurents, and Mariano Carrion-Vazquez. 2018. “Efficient and Simplified Nanomechanical Analysis of Intrinsically Disordered Proteins.” Nanoscale 10 (35): 16857–67. https://doi.org/10.1039/c8nr02785d.

Ferreon, Allan Chris M, Crystal R Moran, Yann Gambin, and Ashok A Deniz. 2010. “Single-Molecule Fluorescence Studies of Intrinsically Disordered Proteins.” Methods in Enzymology 472: 179–204. https://doi.org/10.1016/S0076-6879(10)72010-3.

Finn, Robert D, Penelope Coggill, Ruth Y Eberhardt, Sean R Eddy, Jaina Mistry, Alex L Mitchell, Simon C Potter, et al. 2016. “The Pfam Protein Families Database: Towards a More Sustainable Future.” Nucleic Acids Research 44 (D1): D279–85. https://doi.org/10.1093/nar/gkv1344.

Fioriti, Luana, Cory Myers, Yan-You Huang, Xiang Li, Joseph S Stephan, Pierre Trifilieff, Luca Colnaghi, et al. 2015. “The Persistence of Hippocampal-Based Memory Requires Protein Synthesis Mediated by the Prion-like Protein CPEB3.” Neuron 86 (6): 1433–48. https://doi.org/10.1016/j.neuron.2015.05.021.

Fiumara, Ferdinando, Luana Fioriti, Eric R Kandel, and Wayne A Hendrickson. 2010. “Essential Role of Coiled Coils for Aggregation and Activity of Q/N-Rich Prions and PolyQ Proteins.” Cell 143 (7): 1121–35. https://doi.org/10.1016/j.cell.2010.11.042.

Florin, E. L., M. Rief, H. Lehmann, M. Ludwig, C. Dornmair, V. T. Moy, and H. E. Gaub. 1995. “Sensing Specific Molecular Interactions with the Atomic Force Microscope.” Biosensors and Bioelectronics. https://doi.org/10.1016/0956-5663(95)99227-C.

Goktas, Melis, Chuanfu Luo, Ruby May A. Sullan, Ana E. Bergues-Pupo, Reinhard Lipowsky, Ana Vila Verde, and Kerstin G. Blank. 2018. “Molecular Mechanics of Coiled Coils Loaded in the Shear Geometry.” Chemical Science. https://doi.org/10.1039/c8sc01037d.

Goldschmidt, Lukasz, Poh K Teng, Roland Riek, and David Eisenberg. 2010. “Identifying the Amylome, Proteins Capable of Forming Amyloid-like Fibrils.” Proceedings of the National Academy of Sciences of the United States of America 107 (8): 3487–92. https://doi.org/10.1073/pnas.0915166107.

Hake, L E, and J D Richter. 1994. “CPEB Is a Specificity Factor That Mediates Cytoplasmic Polyadenylation during Xenopus Oocyte Maturation.” Cell 79 (4): 617–27. https://doi.org/10.1016/0092-8674(94)90547-9.

Hervas, Ruben, Liying Li, Amitabha Majumdar, Maria Del Carmen Fernandez-Ramirez, Jay R Unruh, Brian D Slaughter, Albert Galera-Prat, et al. 2016. “Molecular Basis of Orb2 Amyloidogenesis and Blockade of Memory Consolidation.” PLoS Biology 14 (1): e1002361. https://doi.org/10.1371/journal.pbio.1002361.

Hervas, Ruben, Javier Oroz, Albert Galera-Prat, Oscar Goni, Alejandro Valbuena, Andres M Vera, Angel Gomez-Sicilia, et al. 2012. “Common Features at the Start of the Neurodegeneration Cascade.” PLoS Biology 10 (5): e1001335. https://doi.org/10.1371/journal.pbio.1001335.

Hervas, Ruben, Michael J Rau, Younshim Park, Wenjuan Zhang, Alexey G Murzin, James A J Fitzpatrick, Sjors H W Scheres, and Kausik Si. 2020. “Cryo-EM Structure of a Neuronal Functional Amyloid Implicated in Memory Persistence in Drosophila.” Science (New York, N.Y.) 367 (6483): 1230–34. https://doi.org/10.1126/science.aba3526.

Hou, Fajian, Lijun Sun, Hui Zheng, Brian Skaug, Qiu-Xing Jiang, and Zhijian J Chen. 2011. “MAVS Forms Functional Prion-like Aggregates to Activate and Propagate Antiviral Innate Immune Response.” Cell 146 (3): 448–61. https://doi.org/10.1016/j.cell.2011.06.041.

Improta, S, A S Politou, and A Pastore. 1996. “Immunoglobulin-like Modules from Titin I-Band: Extensible Components of Muscle Elasticity.” Structure (London, England □: 1993) 4 (3): 323–37.

Kayed, Rakez, Elizabeth Head, Floyd Sarsoza, Tommy Saing, Carl W Cotman, Mihaela Necula, Lawrence Margol, et al. 2007. “Fibril Specific, Conformation Dependent Antibodies Recognize a Generic Epitope Common to Amyloid Fibrils and Fibrillar Oligomers That Is Absent in Prefibrillar Oligomers.” Molecular Neurodegeneration 2 (September): 18. https://doi.org/10.1186/1750-1326-2-18.

Kayed, Rakez, Elizabeth Head, Jennifer L Thompson, Theresa M McIntire, Saskia C Milton, Carl W Cotman, and Charles G Glabe. 2003. “Common Structure of Soluble Amyloid Oligomers Implies Common Mechanism of Pathogenesis.” Science (New York, N.Y.) 300 (5618): 486–89. https://doi.org/10.1126/science.1079469.

Keleman, Krystyna, Sebastian Kruttner, Mattias Alenius, and Barry J Dickson. 2007. “Function of the Drosophila CPEB Protein Orb2 in Long-Term Courtship Memory.” Nature Neuroscience 10 (12): 1587–93. https://doi.org/10.1038/nn1996.

Khan, Mohammed Repon, Liying Li, Consuelo Perez-Sanchez, Anita Saraf, Laurence Florens, Brian D Slaughter, Jay R Unruh, and Kausik Si. 2015. “Amyloidogenic Oligomerization Transforms Drosophila Orb2 from a Translation Repressor to an Activator.” Cell 163 (6): 1468–83. https://doi.org/10.1016/j.cell.2015.11.020.

Krishnan, Rajaraman, Jessica L Goodman, Samrat Mukhopadhyay, Chris D Pacheco, Edward A Lemke, Ashok A Deniz, and Susan Lindquist. 2012. “Conserved Features of Intermediates in Amyloid Assembly Determine Their Benign or Toxic States.” Proceedings of the National Academy of Sciences of the United States of America 109 (28): 11172–77. https://doi.org/10.1073/pnas.1209527109.

Kruttner, Sebastian, Barbara Stepien, Jasprina N Noordermeer, Mieke A Mommaas, Karl Mechtler, Barry J Dickson, and Krystyna Keleman. 2012. “Drosophila CPEB Orb2A Mediates Memory Independent of Its RNA-Binding Domain.” Neuron 76 (2): 383–95. https://doi.org/10.1016/j.neuron.2012.08.028.

Kruttner, Sebastian, Lisa Traunmuller, Ugur Dag, Katharina Jandrasits, Barbara Stepien, Nirmala Iyer, Lee G Fradkin, Jasprina N Noordermeer, Brett D Mensh, and Krystyna Keleman. 2015. “Synaptic Orb2A Bridges Memory Acquisition and Late Memory Consolidation in Drosophila.” Cell Reports 11 (12): 1953–65. https://doi.org/10.1016/j.celrep.2015.05.037.

Li, Liying, Consuelo Perez Sanchez, Brian D Slaughter, Yubai Zhao, Mohammed Repon Khan, Jay R Unruh, Boris Rubinstein, and Kausik Si. 2016. “A Putative Biochemical Engram of Long-Term Memory.” Current Biology□: CB 26 (23): 3143–56. https://doi.org/10.1016/j.cub.2016.09.054.

Lilliu, Elena, Veronica Villeri, Ilaria Pelassa, Federico Cesano, Domenica Scarano, and Ferdinando Fiumara. 2018. “Polyserine Repeats Promote Coiled Coil-Mediated Fibril Formation and Length-Dependent Protein Aggregation.” Journal of Structural Biology 204 (3): 572–84. https://doi.org/10.1016/j.jsb.2018.09.001.

Liu, Cong, Minglei Zhao, Lin Jiang, Pin-Nan Cheng, Jiyong Park, Michael R Sawaya, Anna Pensalfini, et al. 2012. “Out-of-Register Beta-Sheets Suggest a Pathway to Toxic Amyloid Aggregates.” Proceedings of the National Academy of Sciences of the United States of America 109 (51): 20913–18. https://doi.org/10.1073/pnas.1218792109.

Lopez-Alonso, Jorge Pedro, Marta Bruix, Josep Font, Marc Ribo, Maria Vilanova, Maria Angeles Jimenez, Jorge Santoro, Carlos Gonzalez, and Douglas V Laurents. 2010. “NMR Spectroscopy Reveals That RNase A Is Chiefly Denatured in 40% Acetic Acid: Implications for Oligomer Formation by 3D Domain Swapping.” Journal of the American Chemical Society 132 (5): 1621–30. https://doi.org/10.1021/ja9081638.

Lupas, A, M Van Dyke, and J Stock. 1991. “Predicting Coiled Coils from Protein Sequences.” Science (New York, N.Y.) 252 (5009): 1162–64. https://doi.org/10.1126/science.252.5009.1162.

Majumdar, Amitabha, Wanda Colon Cesario, Erica White-Grindley, Huoqing Jiang, Fengzhen Ren, Mohammed Repon Khan, Liying Li, et al. 2012. “Critical Role of Amyloid-like Oligomers of Drosophila Orb2 in the Persistence of Memory.” Cell 148 (3): 515–29. https://doi.org/10.1016/j.cell.2012.01.004.

Mitchell, Alex, Hsin-Yu Chang, Louise Daugherty, Matthew Fraser, Sarah Hunter, Rodrigo Lopez, Craig McAnulla, et al. 2015. “The InterPro Protein Families Database: The Classification Resource after 15 Years.” Nucleic Acids Research 43 (Database issue): D213–21. https://doi.org/10.1093/nar/gku1243.

Mukhopadhyay, Samrat, Rajaraman Krishnan, Edward A Lemke, Susan Lindquist, and Ashok A Deniz. 2007. “A Natively Unfolded Yeast Prion Monomer Adopts an Ensemble of Collapsed and Rapidly Fluctuating Structures.” Proceedings of the National Academy of Sciences of the United States of America 104 (8): 2649–54. https://doi.org/10.1073/pnas.0611503104.

Nagai, Yoshitaka, Takashi Inui, H Akiko Popiel, Nobuhiro Fujikake, Kazuhiro Hasegawa, Yoshihiro Urade, Yuji Goto, Hironobu Naiki, and Tatsushi Toda. 2007. “A Toxic Monomeric Conformer of the Polyglutamine Protein.” Nature Structural & Molecular Biology 14 (4): 332–40. https://doi.org/10.1038/nsmb1215.

Oberhauser, Andres F, and Mariano Carrion-Vazquez. 2008. “Mechanical Biochemistry of Proteins One Molecule at a Time.” The Journal of Biological Chemistry 283 (11): 6617–21. https://doi.org/10.1074/jbc.R700050200.

Oroz, Javier, Ruben Hervas, and Mariano Carrion-Vazquez. 2012. “Unequivocal Single-Molecule Force Spectroscopy of Proteins by AFM Using PFS Vectors.” Biophysical Journal 102 (3): 682–90. https://doi.org/10.1016/j.bpj.2011.12.019.

Piotto, M, V Saudek, and V Sklenar. 1992. “Gradient-Tailored Excitation for Single-Quantum NMR Spectroscopy of Aqueous Solutions.” Journal of Biomolecular NMR 2 (6): 661–65.

Ramos-Martin, Francisco, Ruben Hervas, Mariano Carrion-Vazquez, and Douglas V Laurents. 2014. “NMR Spectroscopy Reveals a Preferred Conformation with a Defined Hydrophobic Cluster for Polyglutamine Binding Peptide 1.” Archives of Biochemistry and Biophysics 558 (September): 104–10. https://doi.org/10.1016/j.abb.2014.06.025.

Raveendra, Bindu L, Ansgar B Siemer, Sathyanarayanan V Puthanveettil, Wayne A Hendrickson, Eric R Kandel, and Ann E McDermott. 2013. “Characterization of Prion-like Conformational Changes of the Neuronal Isoform of Aplysia CPEB.” Nature Structural & Molecular Biology 20 (4): 495–501. https://doi.org/10.1038/nsmb.2503.

Schwaiger, Ingo, Clara Sattler, Daniel R Hostetter, and Matthias Rief. 2002. “The Myosin Coiled-Coil Is a Truly Elastic Protein Structure.” Nature Materials 1 (4): 232–35. https://doi.org/10.1038/nmat776.

Si, Kausik, Yun-Beom Choi, Erica White-Grindley, Amitabha Majumdar, and Eric R Kandel. 2010. “Aplysia CPEB Can Form Prion-like Multimers in Sensory Neurons That Contribute to Long-Term Facilitation.” Cell 140 (3): 421–35. https://doi.org/10.1016/j.cell.2010.01.008.

Si, Kausik, Susan Lindquist, and Eric R Kandel. 2003. “A Neuronal Isoform of the Aplysia CPEB Has Prion-like Properties.” Cell 115 (7): 879–91. https://doi.org/10.1016/s0092-8674(03)01020-1.

Stephan, Joseph S, Luana Fioriti, Nayan Lamba, Luca Colnaghi, Kevin Karl, Irina L Derkatch, and Eric R Kandel. 2015. “The CPEB3 Protein Is a Functional Prion That Interacts with the Actin Cytoskeleton.” Cell Reports 11 (11): 1772–85. https://doi.org/10.1016/j.celrep.2015.04.060.

Tomita, Kenji, H Akiko Popiel, Yoshitaka Nagai, Tatsushi Toda, Yuji Yoshimitsu, Hiroaki Ohno, Shinya Oishi, and Nobutaka Fujii. 2009. “Structure-Activity Relationship Study on Polyglutamine Binding Peptide QBP1.” Bioorganic & Medicinal Chemistry 17 (3): 1259–63. https://doi.org/10.1016/j.bmc.2008.12.018.

Valbuena, Alejandro, Javier Oroz, Ruben Hervas, Andres Manuel Vera, David Rodriguez, Margarita Menendez, Joanna I Sulkowska, Marek Cieplak, and Mariano Carrion-Vazquez. 2009. “On the Remarkable Mechanostability of Scaffoldins and the Mechanical Clamp Motif.” Proceedings of the National Academy of Sciences of the United States of America 106 (33): 13791–96. https://doi.org/10.1073/pnas.0813093106.

Volkov, Kirill V, Anna Yu Aksenova, Malle J Soom, Kirill V Osipov, Anton V Svitin, Cornelia Kurischko, Irina S Shkundina, Michael D Ter-Avanesyan, Sergey G Inge-Vechtomov, and Ludmila N Mironova. 2002. “Novel Non-Mendelian Determinant Involved in the Control of Translation Accuracy in Saccharomyces Cerevisiae.” Genetics 160 (1): 25–36.

Wickner, R B. 1994. “[URE3] as an Altered URE2 Protein: Evidence for a Prion Analog in Saccharomyces Cerevisiae.” Science (New York, N.Y.) 264 (5158): 566–69. https://doi.org/10.1126/science.7909170.

Wurth, Christine, Nathalie K Guimard, and Michael H Hecht. 2002. “Mutations That Reduce Aggregation of the Alzheimer’s Abeta42 Peptide: An Unbiased Search for the Sequence Determinants of Abeta Amyloidogenesis.” Journal of Molecular Biology 319 (5): 1279–90. https://doi.org/10.1016/S0022-2836(02)00399-6.

Xue, Bin, Roland L Dunbrack, Robert W Williams, A Keith Dunker, and Vladimir N Uversky. 2010. “PONDR-FIT: A Meta-Predictor of Intrinsically Disordered Amino Acids.” Biochimica et Biophysica Acta 1804 (4): 996–1010. https://doi.org/10.1016/j.bbapap.2010.01.011.

