## Supplemental Material for "Divergent CPEB prion-like domains reveal different assembly mechanisms for a generic amyloid-like fold"

**Supporting Figures**

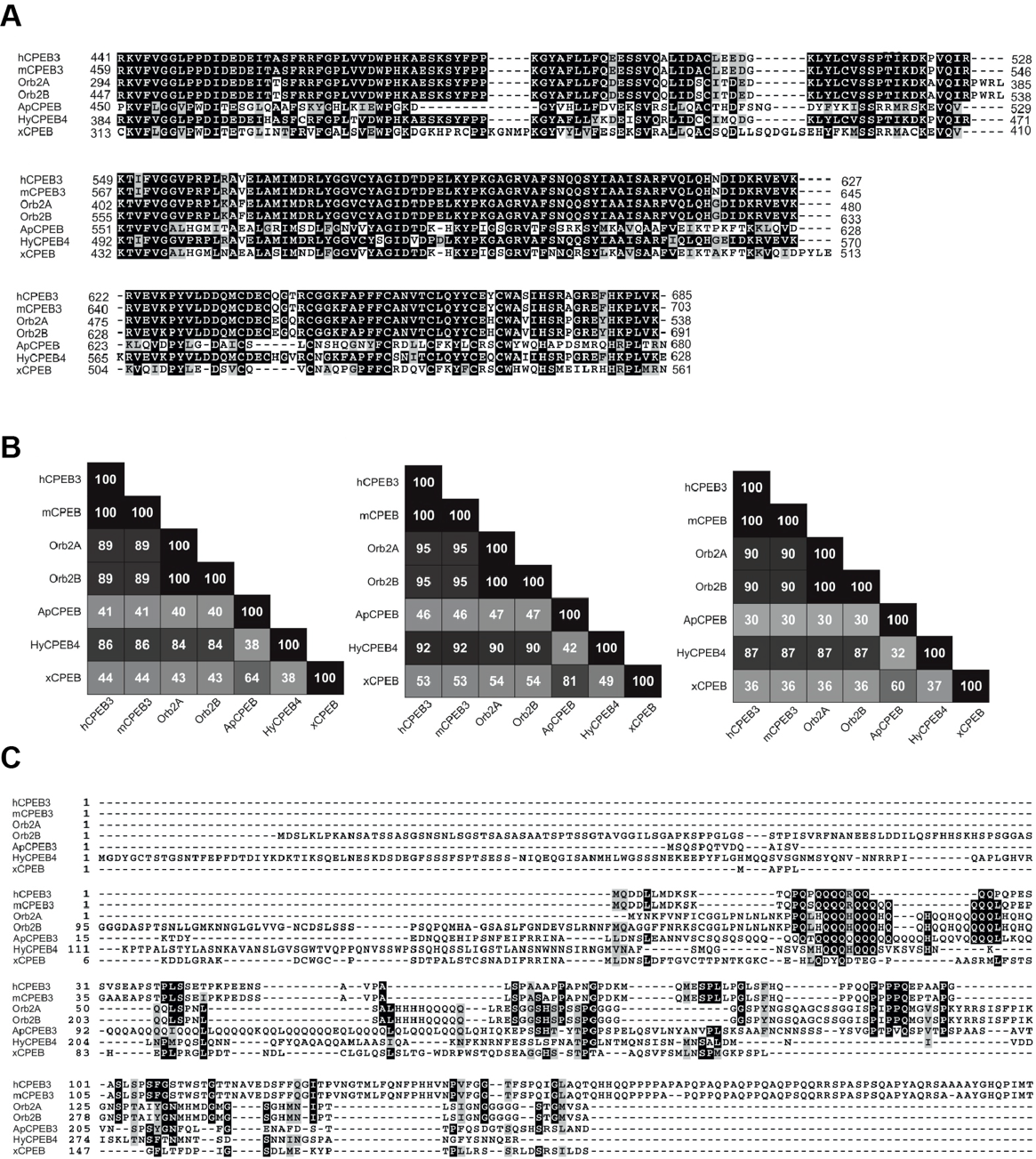

**Figure S1. Sequence analysis of CPEB proteins. A)** Sequence aligments of the structured C-terminal domains RRM1 (top panel), RRM2 (middle) and ZZ (bottom). **B)** Pairwise sequence identity represented as a matrix: RRM1 (left panel), RRM2 (middle) and ZZ (right). **C)** Sequence comparison of the N-terminal (disordered) regions. The sequence considered corresponds to the fragment from the N-terminus to the first residue of the RRM1 motif. This region is much less conserved than the C-terminal domains.

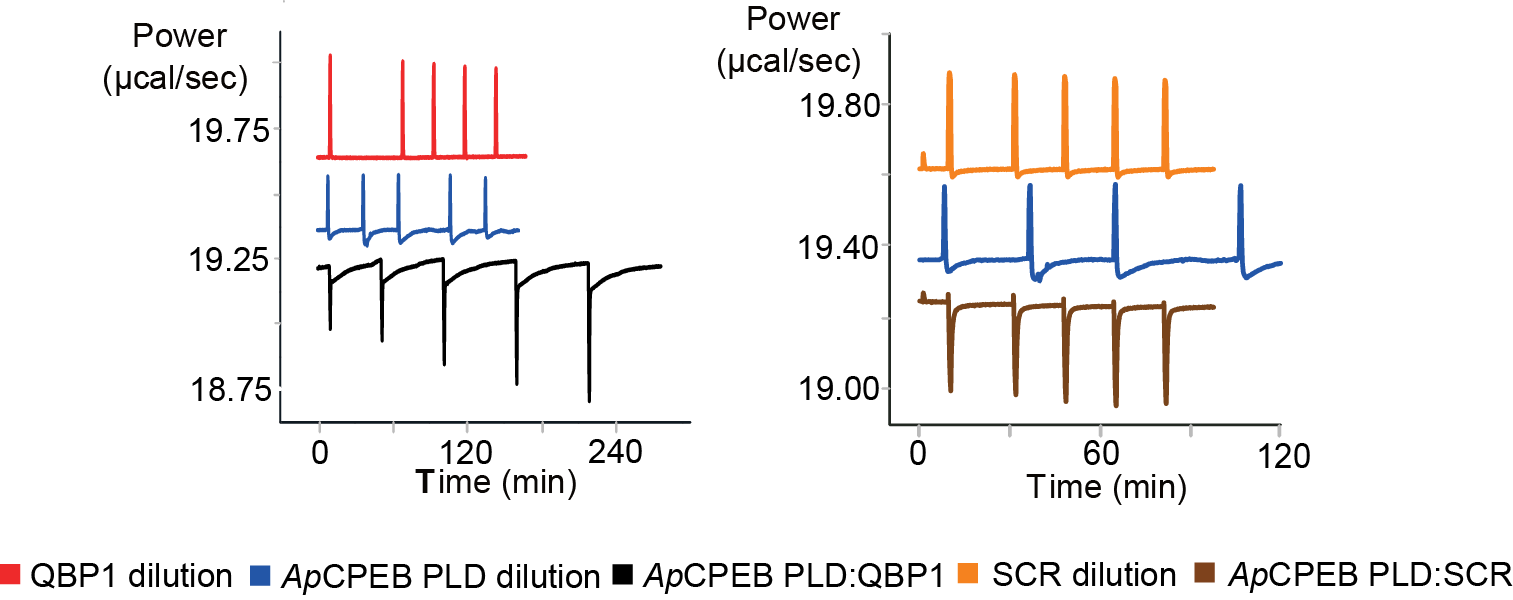

**Figure S2. Interaction of QBP1 with *Ap*CPEB PLD.** Representative calorimetric ITC traces for the injection of 550 μM *Ap*CPEB PLD (10-25 *µ*l each) into 75 μM QBP1 (left panel) or 51 μM SCR (right panel) peptides. Measurements were performed in PBS pH 7.4, at 25 ˚C. Traces in black and brown correspond to the heat released upon injection of *Ap*CPEB PLD into the ITC cell loaded with an excess of QBP1 or SRC, respectively. Blue, red and orange traces correspond to the *Ap*CPEB PLD, QBP1, and SRC dilutions, respectively.

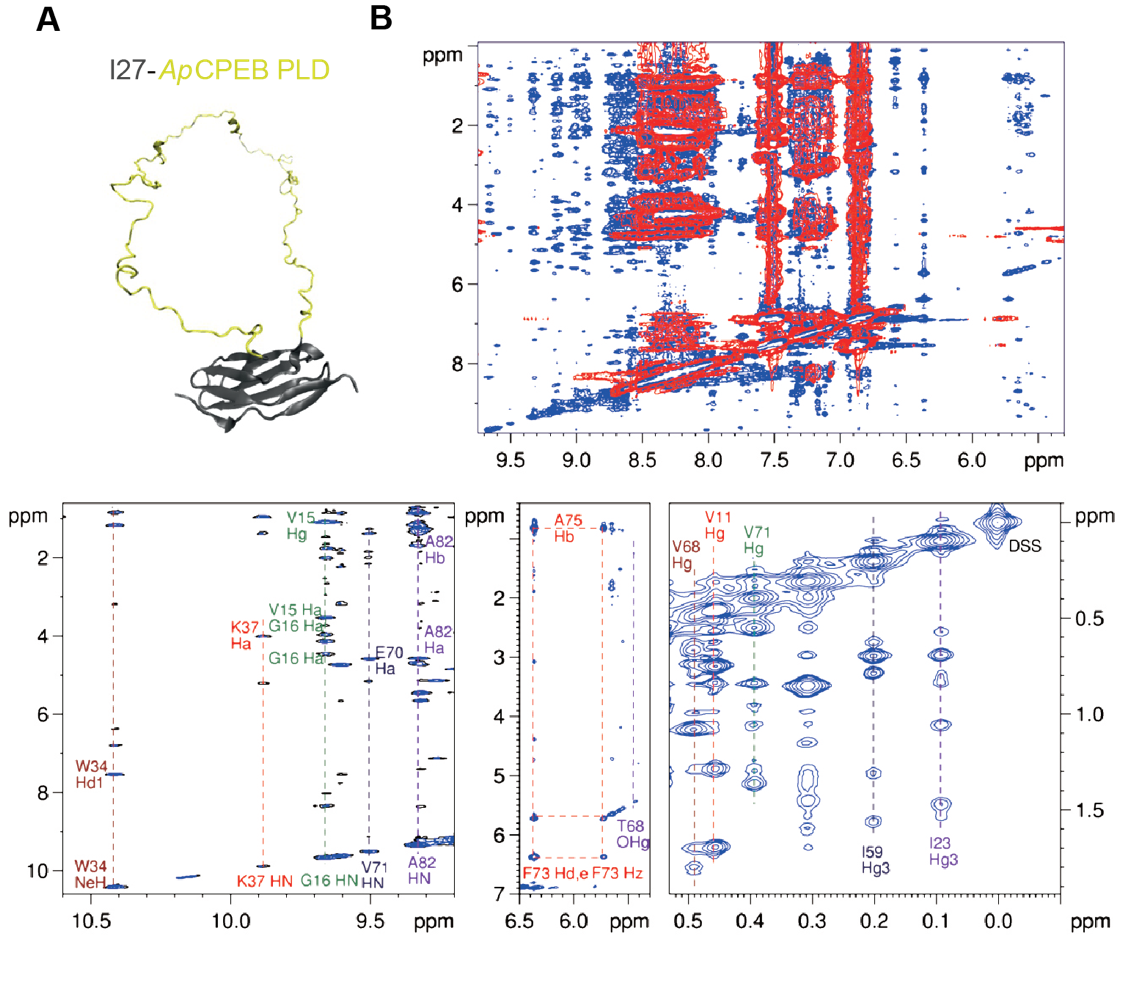

**Figure S3. Validation of the mechanical protection strategy used in AFM-SMFS experiments**. **A)** Schematic cartoon representation of I27-*Ap*CPEB PLD carrier-guest construction colored in gray and yellow, respectively. The representation was prepared with MODELLER and displayed by Visual Molecular Dynamics v1.8.6, using atomic coordinates for titin I27 (PDB code 1TIT) and off-template modelling for *Ap*CPEB PLD. **B)** 2D ^1^H NOESY spectra of *Ap*CPEB PLD (red) and I27-*Ap*CPEB PLD (blue). The upper panel shows the downfield region of the spectra. Whereas I27-*Ap*CPEB PLD gives rise to many signals with diverse range of chemical shifts, those of *Ap*CPEB PLD are clumped together in regions whose chemical shift is typical of disordered peptides and proteins (Lopez-Alonso et al. 2010). Additional 3D heteronuclear spectra would be necessary to detect small content of *β-* or *α-*structures. The bottom panels show selected local regions of the fusion protein spectrum and some peaks which are unambiguously assigned to ^1^H in I27 labeled, which is conclusive evidence of the I27 native structure conservation.

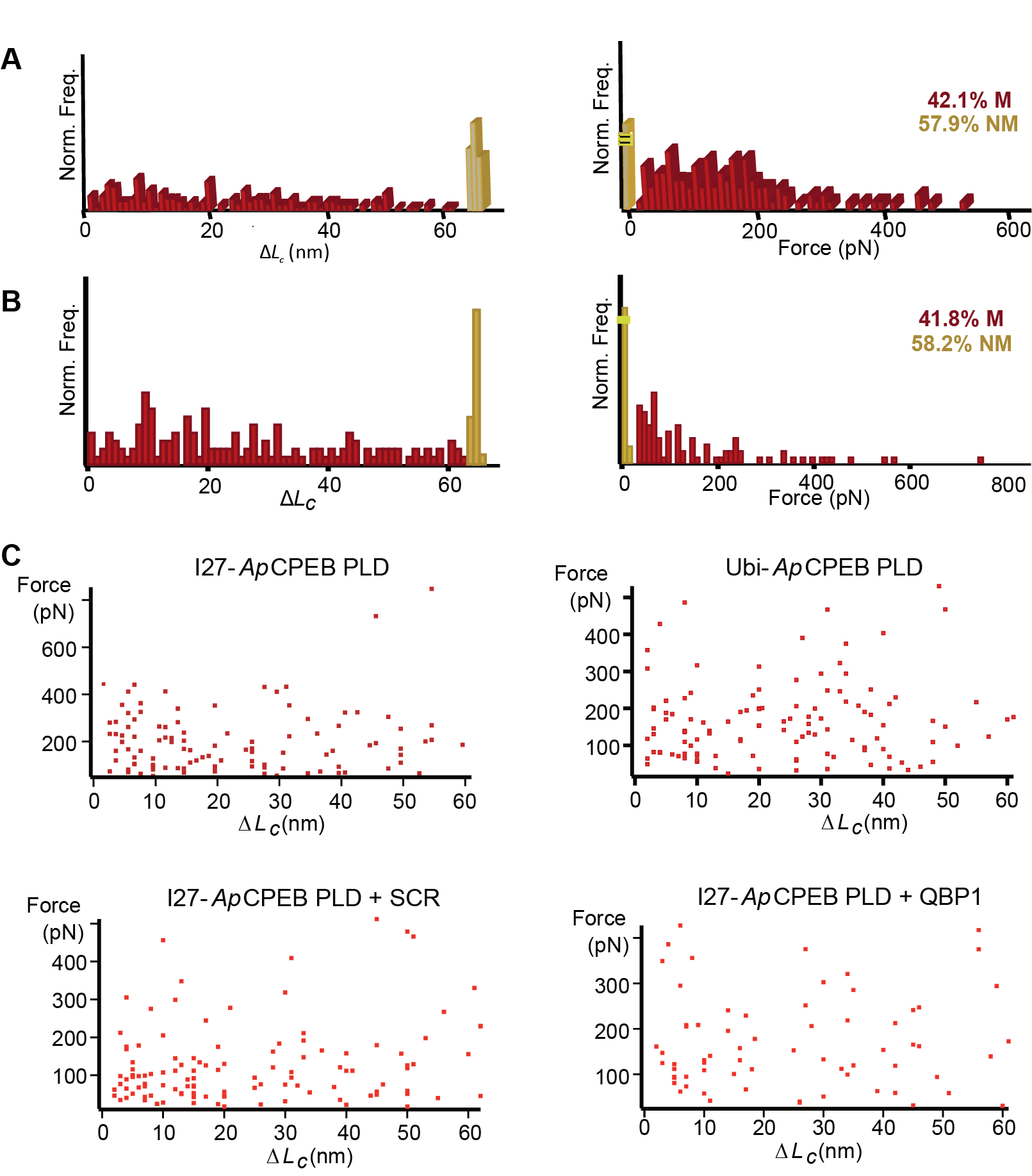

**Figure S4. Additional data from SMFS experiments. A)** Δ*L_c_* (left panel) and *F*_u_ (right panel) histograms from pFS-2-*Ap*CPEB PLD using Ubi as a carrier (*n*=155). **B)** Effect of DMSO in the *Ap*CPEB PLD SMFS analysis. Δ*L_c_* (left panel) and *F*_u_ (right panel) histograms from experiments in presence of 0.01 % DMSO show that this concentration, which is used to dissolve QBP1 and SCR, does not perturb the distributions (*n*=97). **C)** Scatter plots show that, in the M conformers of any condition tested, there is no clustering between *F*_u_ and Δ*L_c_*, which suggests that no preferred structures are formed.

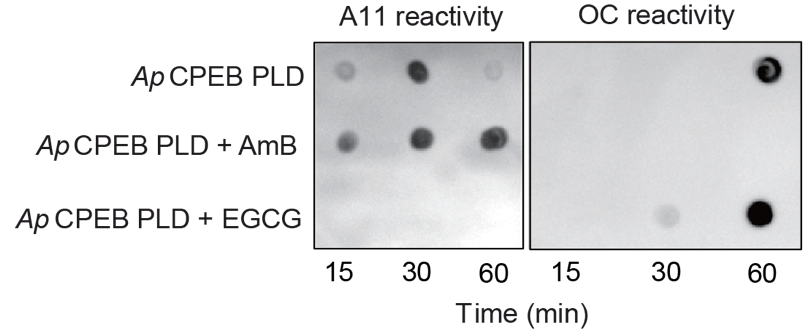

**Figure S5. Different species of *Ap*CPEB PLD are trapped by small molecules.** Immuno-dot blot analysis of the *Ap*CPEB PLD aggregates trapped by AmB or EGCG. Species treated with EGCG interact with OC antibody, whereas species formed in presence of AmB react with A11, but no with OC antibody.

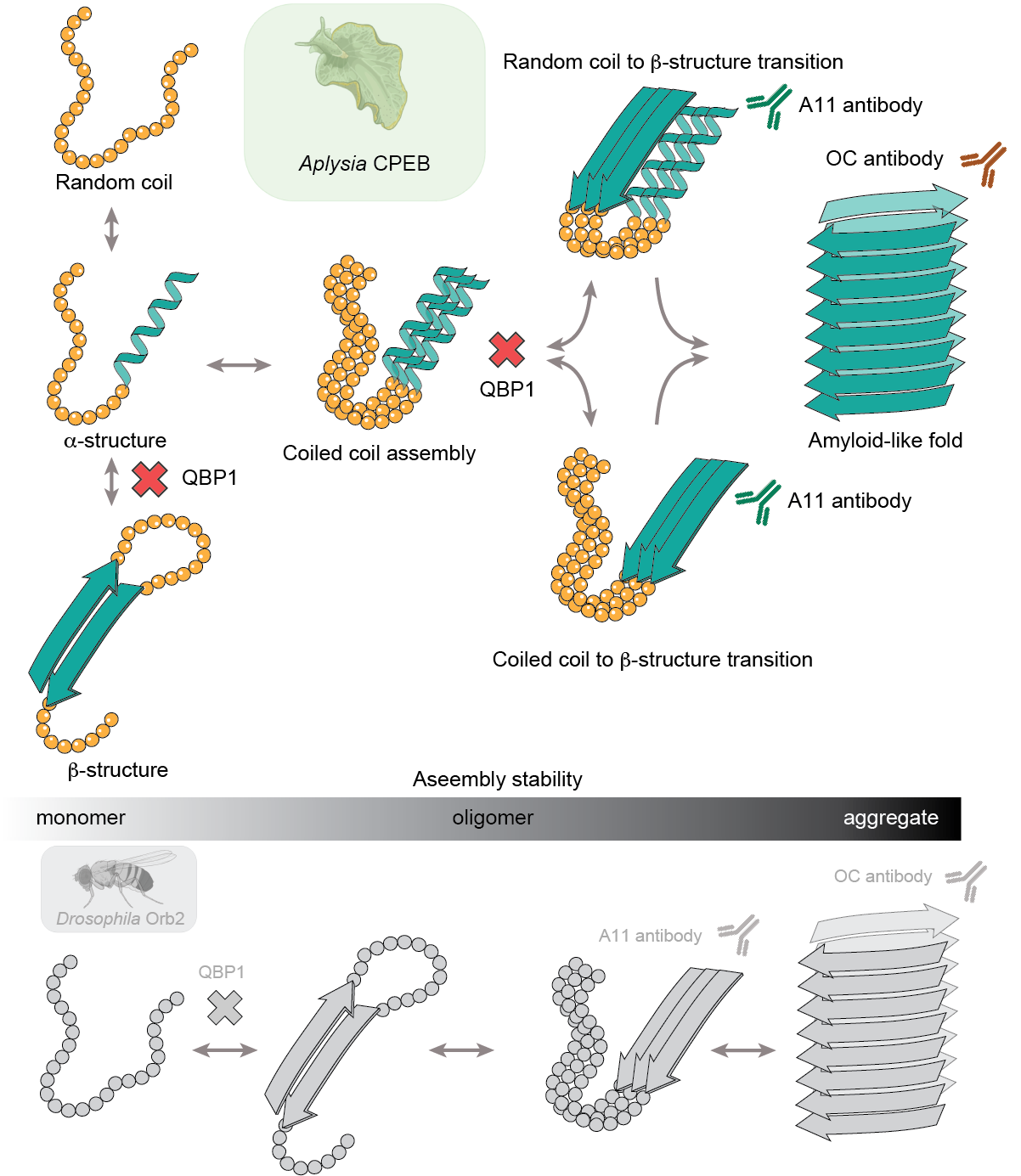

**Figure S6. Possible role of coiled coils in the assembly pathway of *Ap*CPEB PLD**. Monomeric *Ap*CPEB PLD fluctuates among *β*-, *α*- and random coil conformations. In contrast to monomeric Orb2 PLD (bottom row) (Hervas et al. 2016), which is in a random coil conformation, the association of *α*-helices present in monomeric *Ap*CPEB PLD, lead to the formation of CC species. CCs may represent intermediate assembly structures facilitating their own conversion into *β*-sheet multimers. Alternatively, CCs may mediate the conformational transition of neighboring random coil segments into *β*-sheet multimers CC (Fiumara et al. 2010; M. Chen, Zheng, and Wolynes 2016). *Ap*CPEB PLD and Orb2 PLD pathways would converge in the formation of A11-reactive multimers, evolving to the formation of a common OC-reactive, amyloid-like fold. QBP1, known to interact with polyQ segments, would block the monomer transition to a *β*-rich conformer and potentially the CCs transition to *β*-sheet multimers, which would result in a reduction of M events in SMFS and, hence, in the blockade of the amyloid-like assembly.

| **Protein** | **n** | **M (%)** | **NM (%)** | **#peaks/mol.** | ***F vs*. Δ*L_c_*** |
| --- | --- | --- | --- | --- | --- |
| **Ubi-*Ap*CPEB PLD** | 155 | 42.1 | 57.9 | 1.87 | Uncorrelated |
| **I27-*Ap*CPEB PLD** | 145 | 40.0 | 60.0 | 1.72 | Uncorrelated |
| **I27-*Ap*CPEB PLD +QBP1** | 186 | 21.5 | 78.5 | 1.68 | Uncorrelated |
| **I27-*Ap*CPEB PLD +SCR** | 144 | 43.3 | 56.7 | 1.77 | Uncorrelated |
| **I27-*Ap*CPEB PLD +DMSO** | 97 | 41.8 | 58.2 | 1.81 | Uncorrelated |

**Supplemental Table 1. Summary of SMFS analysis of *Ap*CPEB PLD**.

| **Construct** | **Oligonucleotide (sequence 5’ to 3’)** |
| --- | --- |
| ***Ap*CPEB PLD 5’** | CTA*GCTAG*CCATGCAAGCCATGGCCGT |
| ***Ap*CPEB PLD 3’** | CCG*CTCGAG*CTACTATGGAACCAGGCGTGTA |
| **I27-*Ap*CPEB PLD 5’**  **Ubi-*Ap*CPEB PLD 5’**  **pFS2-I27-*Ap*CPEB PLD 5’**  **pFS-2-Ubi-*Ap*CPEB PLD 5’** | CCAA*ACCGGT*ATGCAAGCCATGGCCGT |
| **pFS-2-I27-*Ap*CPEB PLD 3’**  **pFS-2-Ubi-*Ap*CPEB PLD 3’** | TCC*CCCGGG*TGGACCAGGCGTGTA |

**Supplemental Table 2. Summary of the oligonucleotides used.** The restriction sites introduced by PCR into the amplified sequences are highlighted in italics. Underlined sequences correspond to stop codons. The extra sequences added to the end of each restriction site were chosen on the basis of the recommendations from New England Biolabs, which enhance the digestion efficiency of linear DNA sequences. All oligonucleotides used were purchased from Sigma-Aldrich.
